# CytoCellDB: A Resource Database For Classification and Analysis of Extrachromosomal DNA in Cancer

**DOI:** 10.1101/2023.12.18.572197

**Authors:** Jacob Fessler, Stephanie Ting, Hong Yi, Santiago Haase, Jingting Chen, Saygin Gulec, Yue Wang, Nathan Smyers, Kohen Goble, Danielle Cannon, Aarav Mehta, Christina Ford, Elizabeth Brunk

## Abstract

Extrachromosomal DNA (ecDNA), or double minute chromosomes, are established cytogenetic markers for malignancy and genome instability. More recently, the cancer community has gained a heightened awareness of the roles of ecDNA in cancer proliferation, drug resistance and epigenetic remodeling. A current hindrance to understanding the biological roles of ecDNA is the lack of available cell line model systems with experimental cytogenetic data that confirm ecDNA status. Although several recent landmark studies have identified common cell lines and tumor models with ecDNA, the current sample size limits our ability to detect ecDNA-driven molecular differences due to limitations in power. Increasing the number of model systems known to express ecDNA would provide new avenues for understanding the fundamental underpinnings of ecDNA biology and would unlock a wealth of potential targeting strategies for ecDNA-driven cancers. To bridge this gap, we created CytoCellDB, a resource that provides karyotype annotations and leverages publicly available global cell line data from the Cancer Dependency Map (DepMap) and the Cancer Cell Line Encyclopedia (CCLE). Here, we identify 139 cell lines that express ecDNA, which is a 200% increase from the current sample size. We expanded the total number of cancer cell lines with ecDNA annotations to 577, which is a 400% increase or 31% of cell lines in CCLE/ DepMap. We demonstrate that a strength of CytoCellDB is the ability to interrogate ecDNA, and a compendium of other chromosomal aberrations, in the context of cancer-specific vulnerabilities, drug sensitivities, and molecular data (genomics, transcriptomics, methylation, proteomics). We anticipate that CytoCellDB will advance cytogenomics research and population-scale discoveries related to ecDNA as well as provide insights into strategies and best practices for determining novel therapeutics that overcome ecDNA-driven drug resistance.

---

Extrachromosomal DNA (ecDNA) are large (∼kilo to megabase) acentric, atelomeric, circular DNAs that enable cancer cells to amplify key driver genes outside of chromosomes and change how these genes are regulated, replicated and divided. While this sounds like the basis of a fictional biological thriller, ecDNA are more common than expected^1–3^; they occur in as high as 14-20%^2,4^ of tumors from oncology patients and have been established cytogenetic markers for malignancy and hard-to-treat tumors for 40 years^5^ (often called double minute chromosomes). Surprisingly, despite their clinical relevance, it remains largely unknown how frequently they occur in cancer cell line model systems. Some of the most commonly used cancer cell line models have been extensively profiled by Cancer Cell Line Encyclopedia (CCLE)^6^ and Dependency Map^7^ (DepMap) projects, which host large, data-rich, population-scale repositories and provide multi-omic data for nearly 2,000 cancer cell lines. Yet, only a small fraction (5%) of cell lines have been experimentally tested for ecDNA, which exposes a crucial knowledge gap that limits basic cancer research in many ways. Understanding how systems are impacted by ecDNA is an urgent, unmet need that will change the way we analyze genomics and sequencing data, develop drug screens and understand drug resistance.

Extrachromosomal DNA differs from chromosomal DNA in several important ways. First, the genetic material of ecDNA is often composed of genes from multiple chromosomes and differs in genetic sequence and count from cell to cell (*Fig. 1A*). Genetically distinct ecDNA species are visualized using Fluorescence in situ Hybridization (FISH) and Scanning Electron Microscopy (SEM) (*Fig. 1B*). For example, in the gastric cancer cell line, SNU16, at least three distinct species of ecDNA exist independently from chromosomes; those that localize the MYC (chr 8) oncogene, those that localize the FGFR2 (chr 10) oncogene, and those that co-localize MYC (chr 8), FGFR2 and CD44 (chr 11) genes. These unique species lead to novel breakpoints, in which genetic material from two different chromosomes come together (*Fig. 1C*). The breakpoint regions on multimers have unique properties compared to chromosomal DNA in that they introduce new transcriptional regulatory interactions^1^. The circular structure of ecDNA is thought to make genes more accessible to transcription machinery^8^, potentially changing oncogene expression levels. Lastly, cells with ecDNA have heightened levels of cell-population heterogeneity because ecDNA replicates autonomously and divides unevenly into daughter cells, thereby generating single cells with widely varying ecDNA counts (*Fig. 1D*). Extreme cell-to-cell heterogeneity may increase drug resistance by elevating the number of ecDNA in cells^9^, eliminating ecDNA in cells^10^, or reintegrating ecDNA in homogenous staining regions (HSRs) in chromosomes and restoring them upon drug withdrawal^11^. Increasing our knowledge of which cell lines express ecDNA would open new avenues toward understanding the impact of ecDNA on genome function, cancer cell fitness, therapeutic response and drug evasion.

**Figure 1.**
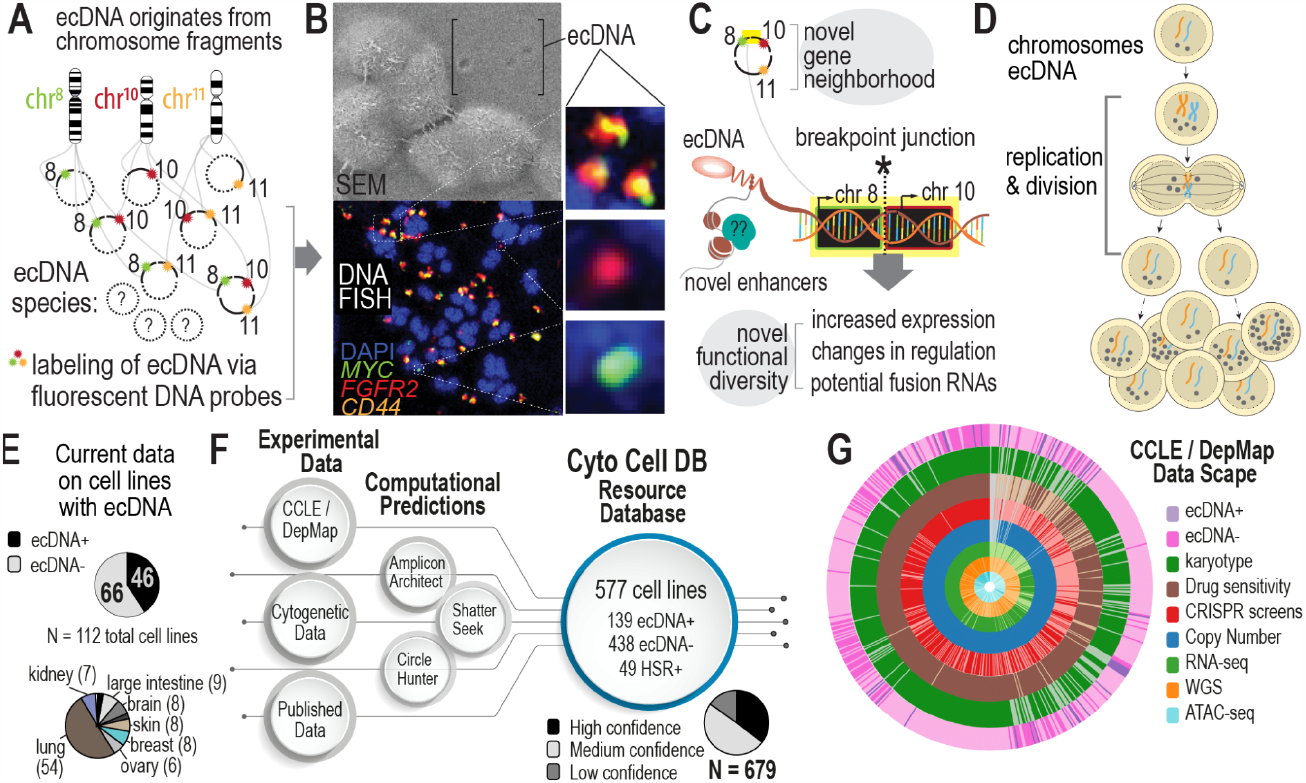
Increasing the scope of ecDNA knowledge in common cancer cell lines. **A**. The biogenesis mechanism for ecDNA is not well understood, but ecDNA are focal amplifications of genomic regions that can multimerize to co-localize on the same ecDNA amplicon. Depending on the cell line, this can lead to many diverse species. **B**. Microscopy methods, such as scanning electron microscopy (SEM) and DNA Fluorescence in situ Hybridization (FISH) are gold standard approaches for visualizing ecDNA. DNA FISH uses fluorescent probes that bind to specific DNA sequences that indicate which regions of the genome are amplified by ecDNA. Multiple species can be observed depending on which fluorescent probe is detected. **C**. Co-localization of genes from different chromosomes on the same ecDNA amplicon may lead to novel biological functionalities. A breakpoint junction is where two parts of different chromosomes come together, potentially producing fusion DNA or may lead to changes in regulation and transcription through enhancer remodeling. **D**. Asymmetric division of ecDNA molecules into daughter cells during replication and division. Multiple rounds of replication leads to a heterogeneous mixture of cells with various ecDNA counts and species. **E**. Recent efforts to probe ecDNA status in cell line populations provides data on 112 cell lines, in which 46 are labeled as “true positive” and 66 are labeled as “true negative.” These cell lines mostly consist of small cell lung cancer (54%). **F**. CytoCellDB represents the largest repository for ecDNA classification, providing queryable data for cell lines, as well as ecDNA and homogenous staining region (HSR) annotations. It also provides computational predictions using several available computational frameworks. Karyotype records were classified as high, medium or low confidence. **G**. CytoCellDB aligns with the Cancer Cell Line Encyclopedia (CCLE) and Dependency Map (DepMap) to enable a multi-omic view of ecDNA biology. A donut plot represents all the available multi-omic data for cell lines with ecDNA annotations.

Identifying more cell lines with ecDNA is hindered by the high costs and low throughput gold standard karyotyping methods (FISH and G-banding). To circumvent these challenges, computational methods have been developed to predict ecDNA expression from next generation sequencing data^12–14^. Algorithms search for circular patterns in contigs from whole genome sequencing (WGS) data or chromatin accessibility (ATAC-seq) data. However, limited numbers (112; *Fig. 1E*) of cell lines with known ecDNA expression hinders the power of computational predictions. What our community needs is a large gold standard dataset that classifies cell line models either as “true positive” (TP; expressing ecDNA) or “true negative” (TN; not expressing ecDNA) using experimental karyotype data. By increasing the sample size of true positive and true negative samples, the power of computational analyses would increase, ensuring validity and practical utility of future analyses of ecDNA.

Here, we present CytoCellDB, a resource database enabling the classification of ecDNA in cancer cell lines. CytoCellDB contains queryable karyotype information for 577 cell lines (31% of CCLE/DepMap), including information on ecDNA, HSRs and other chromosomal aberrations. To our knowledge, CytoCellDB is the largest dataset of its kind, reporting 139 true positive (TP; ecDNA+) samples and 438 true negative (TN; ecDNA-) samples; this represents a 200% increase in total ecDNA+ samples and a 400% in cell lines with ecDNA annotations from current data. We demonstrate that CytoCellDB enables a multi-omic view of ecDNA biology by identifying significant differences in gene expression, gene dependencies and drug sensitivities in ecDNA+ versus ecDNA-cell lines. We use karyotype information to compute the accuracies of two commonly used ecDNA prediction softwares, AmpliconArchitect^13^ and CircleHunter^14^. We develop statistical and machine learning models that predict ecDNA+ and HSR+ from karyotype, RNA and copy number variation data, with accuracies between 78-90% (AUC = .88-.92). We validate three of our model predictions using FISH and G-banding karyotyping experiments. As a resource, CytoCellDB provides new avenues for investigating the molecular impacts of ecDNA on a global scale and will advance the development of treatment strategies, biomarkers, and targeted approaches for cancers driven by ecDNA-based amplifications. CytoCellDB is available online at (http://CytoCellDB.unc.edu).

## Results and Discussion

### Increasing the scope of queryable karyotype data on commonly used cancer cell lines

Several recent efforts^1,2,15^ characterize ecDNA in 112 cancer cell lines, where nearly half of the cell lines are lung cancer model systems (*Fig. 1E*). Of the 112 cell lines with ecDNA classification, 46 are true positives and 66 are true negatives, with an expected frequency of ecDNA expression between 39-44%. Using publicly available karyotype data together with literature curation, we expanded the total number of cancer cell lines with ecDNA annotations to 577, or 31% of cell lines in CCLE/DepMap^6,7^ (*Fig 1F*). Of these, 139 cell lines are annotated as true positives, which is a 200% increase in the number of ecDNA+ model systems. We also annotated 49 cell lines to be HSR+ if they contain homogeneous staining regions (HSR). Finally, 438 cell lines are considered as ecDNA-.

CytoCellDB leverages large-scale, publicly-available multi-omics CCLE/DepMap data to explore underlying biological mechanisms of ecDNA. Publicly available multi-omics data includes WGS, transcriptomics (RNA-seq), copy number variation (CNV), ATAC-seq, drug sensitivity profiles and CRISPR loss-of-function genome-wide screens. For cell lines that have ecDNA+, HSR+ or ecDNA-annotations, 487 have high/medium confidence karyotype data, 457 have RNAseq data, 254 have WGS data, 155 have ATAC-seq data, 507 have small molecule or drug screen data, and 337 have CRISPR-mediated loss-of-function screens (*Fig 1G*).

CytoCellDB represents the most comprehensive cytogenetic resource for cancer cell lines and is unique from other cancer data repositories in several ways. Compared to mutation data repositories, CytoCellDB focuses on karyotype details, which are derived from microscopy experiments that visually detect ecDNA, HSRs, the chromosomal complement, and chromosomal aberrations. To date, there are currently no other databases that provide queryable karyotype details for cancer cell lines nor a database that provides ecDNA annotations from cytogenetics data. Other similar resources include the Mitelman Database^16^, which provides karyotype notes on over 16,000 mixed samples (cell lines and clinical samples), which are not linked to CCLE/DepMap and cannot be mapped to multi-omics data. IGRhCellID^17^ provides raw, uncurated karyotype notes on a smaller pool of CCLE cell lines, but only provides unstructured data (not query-able) and does not include ecDNA or HSR annotations. Finally, recent efforts focusing on ecDNA identification have generated multiple databases^18,19^ that provide computational predictions of ecDNA using available software^13^. To this end, CytoCellDB’s cytogenetic annotations serve as the first and only source of large-scale ground truth data for computational ecDNA predictions and future global analyses on ecDNA+ cell lines.

### Karyotype features significantly differentiate ecDNA+ and ecDNA-cells

We created a pipeline that consists of: (i) manual literature searches for experiments that confirm ecDNA expression, and (ii) algorithm-based mining of karyotype details from the most common cell line vendors (Fig. 2A). First, manual literature review of 72 scientific articles identified 154 cell lines with verified ecDNA+/-status. Second, we developed an html-based text mining algorithm to scrape karyotype details from cell line vendor websites (ATCC and DSMZ). Our algorithm identified records for 533 cell lines; from these, we performed manual curation to extract 22 unique cytogenetic features, (double minute chromosome counts, HSR counts, modal chromosome number, chromosome losses or gains, derivative or marker chromosomes, ploidy classification, percent polyploidy, and others; see *Methods*). We labeled karyotype records as low, medium and highly confident, depending upon the level of details included in the record (see *Methods*) and excluded 102 records that were low confidence. The remaining records combined with our literature search identified 139 ecDNA+ cell lines, 49 HSR+ cell lines, and 438 ecDNA-cell lines. We labeled a cell line ecDNA-if the record specifically stated that the karyotype did not contain ecDNA or HSR (86) or if the record was considered to have medium confidence and did not report ecDNA or HSRs.

**Figure 2.**
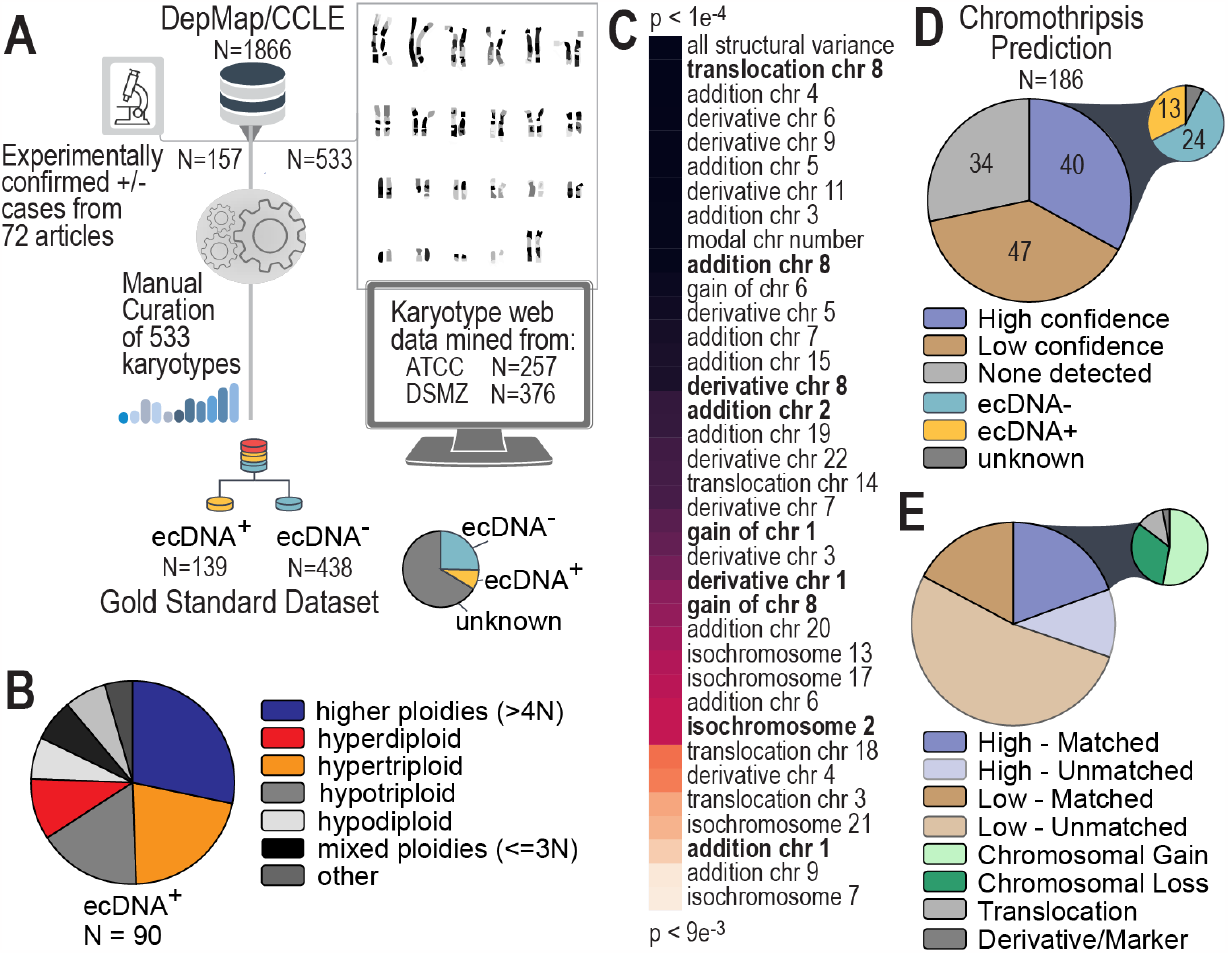
Discovery and Analysis of Hundreds of Cancer Cell Karyotype Data. **A**. We developed a pipeline to mine and organize unstructured karyotype data for hundreds of cancer cell lines. Some records were found in previously published literature while others were taken from cell line vendor websites. All records were manually curated to extract key features. In total, features helped to identify 139 ecDNA+ cell lines, and 438 ecDNA-cell lines, which represents 30% of the cell lines in CCLE/DepMap. **B**. Polyploidy characteristics of ecDNA+ indicate that half the cells have ploidies greater than 3N. **C**. Karyotype features were compared between ecDNA+ and ecDNA-cells to identify the most significant, differentiating properties of ecDNA+ cell lines. **D**. Chromothripsis was predicted in ecDNA+ and ecDNA-cell lines, indicating that genomic rearrangements for many ecDNA+ are not characteristic of chromothripsis. **E**. Alignment of chromosomes involved in chromothripsis predictions (high confidence versus low confidence) with chromosomes observed to have abnormalities (e.g. chromosome gains or losses, translocations or derivative and marker chromosomes) in karyotyping. Matched refers to the percentage of chromosomes predicted to be involved in chromothripsis that were also seen to have aberrations in karyotype data.

We find that key karyotype features differentiate ecDNA+ and ecDNA-cell lines. For example, ecDNA+ cells are enriched in higher polyploidies (greater than 4N, or 92 chromosomes per cell, *Fig. 2B*), which makes up the largest ploidy classification group (29% in ecDNA+ versus 18% ecDNA-cells, respectively). Other karyotype features that significantly differentiate ecDNA+ and ecDNA-cell lines are shown in *Fig. 2C*. Not surprisingly, cells with the largest amount of structural variation were more likely to express ecDNA (p<1e-4 using a Wilcoxon Rank Sum Test for two independent samples). Interestingly, derivative and marker chromosomes involving chromosomes 8, 2 and 1 were more prevalent in cells that express ecDNA; these chromosomes are home to the MYC family of frequently amplified oncogenes, (*MYC, MYC-N* and *MYC-L*), which are known to be extrachromosomally amplified in a number of cell lines^1^. Indeed, significantly high DNA amplifications (copy number > 4, relative to ploidy), found in 76 cell lines, effectively differentiated ecDNA+ (73) and ecDNA-(3) cells. Of these, *MYC* is ranked among the top 5 genes most highly amplified.

Specific patterns of focal amplifications in ecDNA+ cells are thought to arise from chromothripsis–a random, catastrophic shattering of genomic regions, followed by circularization, realignment and loss of a subset of these gene segments^20–22^. We were interested in estimating the frequency of chromothripsis in a larger sample of ecDNA+ and ecDNA-cell lines. Using ShatterSeek^23^, we predicted the likelihood of a chromothripsis event in 184 cell lines (*Fig. 2D*). Remarkably, we find that ecDNA+ cells were not enriched in high confidence predictions (13 of 40 cell lines were ecDNA+). Similarly, ecDNA+ cells were not enriched in low confidence predictions (24 of 47 cell lines were ecDNA+). Strikingly, chromosomes predicted to be involved in chromothripsis are usually chromosomes reported to have aberrations in karyotype data (*Fig. 2E*); 22 of the 34 high confidence chromothripsis predictions involve chromosomes that undergo losses, gains, translocations or have derivative chromosomes.

### EcDNA+ cells exhibit key pathway activities, gene dependencies and drug sensitivities

An exciting application of CytoCellDB is the ability to leverage large-scale multi-omics data from CCLE and DepMap to investigate how ecDNA impacts genome and cellular functioning. One hypothesis is that cells use ecDNA to amplify oncogenes at higher levels than what is possible on chromosomal DNA^8,24,25^. Yet, little attention has been given to understand whether there are genome regions more commonly amplified by ecDNA and whether gene dosage directly impacts gene expression, gene dependencies and drug sensitivities. To investigate these relationships, we developed a pipeline to integrate multi-omics data from CCLE/DepMap^6,7^, the Cancer Therapeutic Response Portal (CTRP) and the Genomics of Drug Sensitivity in Cancer (GDSC) Database^26^ (*Fig. 3A*).

**Figure 3.**
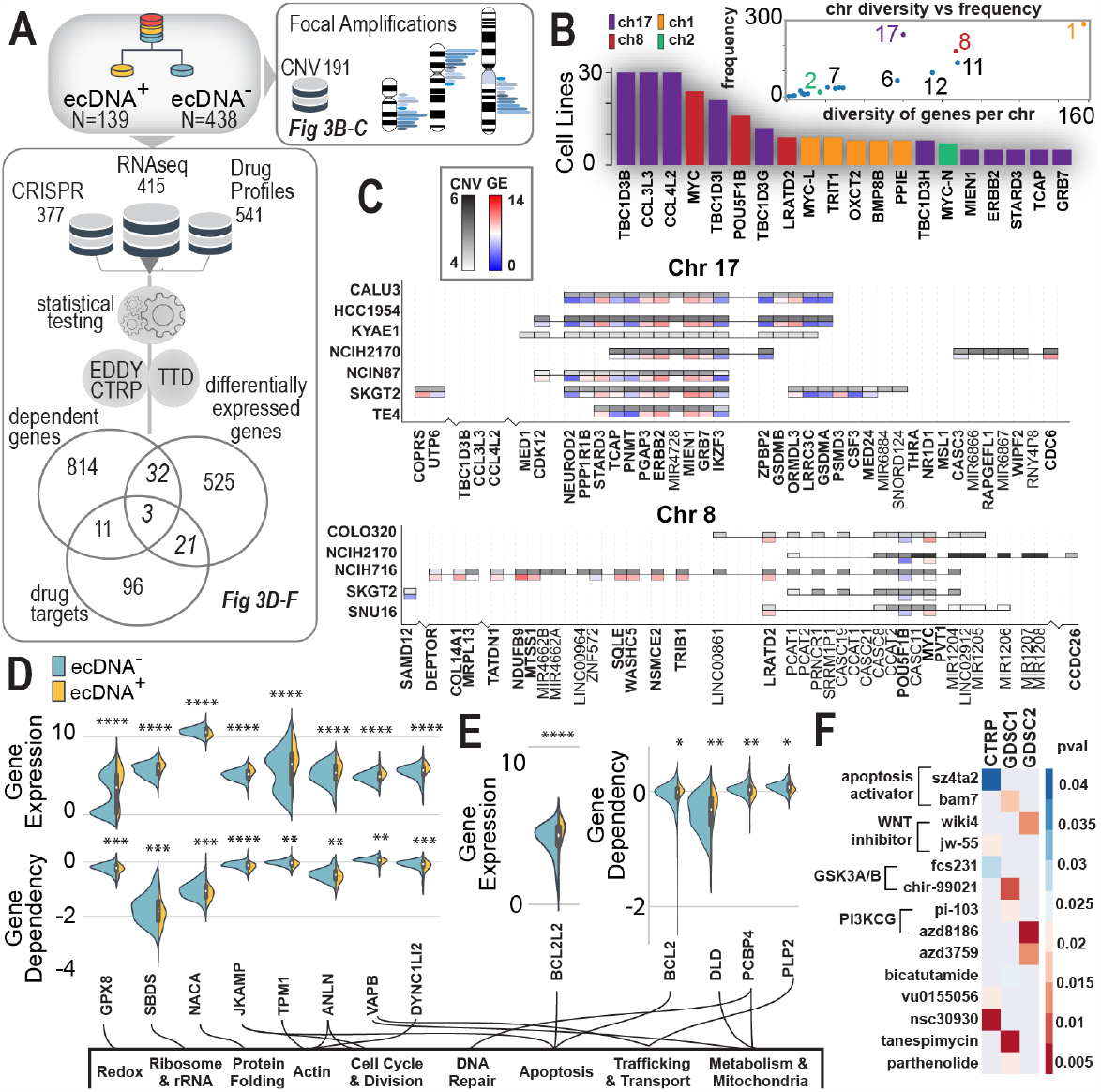
EcDNA+ cells have distinct CNV patterns, pathways, and drug responses. **A**. Multi-omic pipeline that integrates DepMap CRISPR data with DepMap/CCLE transcriptomics data and CTRP/GDSC drug sensitivity profiles to explore uniquely different genes, vulnerabilities and drug responses in ecDNA+ and ecDNA-cell lines. **B**. In ecDNA+ cell lines, certain chromosomes are more commonly amplified (chr 17, 8, 1, and 2) and based on the chromosome, may have a wide range of genes amplified (chr 1) or may have specific regions of the chromosome commonly amplified (chr 2). **C**. Two commonly amplified chromosomal regions in ecDNA+/HSR+ cell lines are chr 8 (bottom) and chr 17 (top). Plotted are heatmaps of CNV and gene expression in these amplified regions, showing patterns of expressions across cell lines. **D**. Comparing differentially expressed genes and genes with significantly different gene dependency scores in ecDNA+ versus ecDNA-cell lines. **E**. Comparing differentially expressed genes and genes involved in drug response, we identified apoptosis (BCL2-family) genes that have significantly higher expression in ecDNA+ cells. **F**. Comparing gene dependencies and genes involved in drug response, we identified three genes in which ecDNA+ cells have lower dependency scores and increased drug sensitivity. For example, a BCL2-family gene (BCL2) has significantly lower dependencies in ecDNA+ cells compared to ecDNA-cells and is a mediator for apoptosis activating drugs, such as sz4ta2 and bam7. In both of these drugs, ecDNA+ cell lines exhibit significantly more sensitive responses compared to ecDNA-cell lines.

We analyzed DNA copy number variation data across 1372 cell lines to understand whether certain genomic regions were more often amplified in ecDNA+ cells. We find that, for large focal amplifications (>10 genes), high density regions on chromosomes 1, 2, 8 and 17 are most commonly found in ecDNA+ cell lines (*Fig. 3B*). We compared the frequency of chromosomal amplification to the number of unique genes originating from a given chromosome (*Fig. 3B*, inset); we find that chromosome 1 is the most diversely amplified chromosome compared to chromosome 2, which is the least diverse, suggesting that only a specific region (containing *MYC-N*) is amplified across cells. While chromosome 17 is the most highly amplified chromosome among ecDNA+ cell lines, *ERBB2*, one of the major oncogenes, is only amplified by a few ecDNA+ cell lines. These highly amplified genomic regions also, generally, had higher RNA levels (Fig. 3C). We observe a common pattern of gene expression in key genes in ecDNA+ cell lines. For example, similar genes, *ERBB2, MIEN1*, and *GRB7*, have higher expression levels, compared to lowly expressed genes, *IKZF3, TCAP, PMNT* and *ZPBP2*; on chromosome 8, *MYC* is heterogeneously expressed across ecDNA+ cell lines.

Identifying differentially expressed genes or gene dependencies in ecDNA+ versus ecDNA-cell lines provides new avenues for drug targeting. We integrated data from gene expression and gene dependencies (from CRISPR-mediated genetic knockdown screens^7,27^) with CytoCellDB. Our search identified 32 genes that are both significantly differentially expressed and have significantly different gene dependency scores (p<0.01 using a Wilcoxon Rank Sum Test for two independent samples). Genes with both higher expression levels and lower dependency scores represent a set of molecular vulnerabilities that are potential drug target candidates (*Fig. 3D*). A gene set enrichment analysis using KEGG (Kyoto Encyclopedia of Genes and Genomes^28^) identified several representative molecular pathways, including cell cycle, immune regulation, DNA repair, oxidative stress, ribosome biogenesis, protein synthesis and degradation, metabolism, apoptosis, endocytosis, intracellular trafficking, cell adhesion and extracellular matrix (ECM) interaction (*Fig. 3D-E*; *Supp Fig. 1*).

Identifying differential drug responses between ecDNA+ and ecDNA-cell lines provides insight into molecular vulnerabilities. We integrated drug sensitivity screens (from CTRP and GDSC), Therapeutic Target Database (TTD)^29^ and EDDY (Evaluation of Differential DependencY)^30,31^ with CytoCellDB to identify and annotate targets and drug mediators that differentiate ecDNA+ and ecDNA-cell lines. The genetic knockdown of several genes (*BCL2, PCBP4*, and *PLP2)* significantly change cell viability in ecDNA+ cell lines; these three genes are known targets or mediators of drugs that also elicit an increase sensitivities in ecDNA+ cells (sz4ta2, bam7, nsc30930; *Fig. 3E*). A family of drugs that elicit a more sensitive response in ecDNA+ cells are apoptosis activators, where multiple genes within the BCL-2 family are also significantly more highly expressed in ecDNA+ cells.

### CytoCellDB enables computational predictions of ecDNA to increase in power

Several computational algorithms have recently been developed to predict ecDNA expression using WGS^12,13,32^ data as well as open chromatin data (ATAC-seq)^14^. Being able to predict the presence of ecDNA through sequencing marks a huge advancement in the field, because gold standard methods for ecDNA detection, (i.e. DNA FISH, G-banding), are low-throughput, cost prohibitive and challenging to scale up. A major unresolved challenge for ecDNA prediction is assessing the accuracy of prediction methods, given the low number of true positives (46 TP; ecDNA+) and true negatives (66 TN; ecDNA-). When sample sizes are too low, prediction methods are underpowered.

CytoCellDB increases the sample sizes of TP samples by 200% and TP+TN samples by 400%, thereby improving prediction power. Landmark studies validated ecDNA expression in NCI60^15,33^ cell lines (60), lung cancer^1^ cell lines (43) and other cell lines^2^ (9). Yet, the number of TP and TN samples are still below the desired numbers to ensure adequate power. For example, if the desired accuracy of predicting ecDNA is 80%, a power analysis reports the minimum sample size should be 97 TP and 97 TN samples (using significance levels for alpha, type I error, and beta, type II error, of 5% and 80%, respectively, and an effect size of 0.4, using Cohen’s h for Proportions^34^; see *Methods*), CytoCellDB provides 139 TP and 438 TN samples (of which 86 are highly confident).

We predicted ecDNA status for 448 and 233 cell lines, using AmpliconArchitect^13^ and CircleHunter^14^, respectively. We find that predictions generated by the software (with default parameters) are consistent with published CircleHunter predictions^14^ (108) and Amplicon Repository^13^ (187). Using CytoCellDB, we contribute ecDNA+ predictions for an additional 189 cell lines that have not yet been analyzed. We compared all predictions to CytoCellDB karyotype annotations to gauge accuracy and precision of computational approaches (*Fig. 4A*). For AmpliconArchitect, we report accuracies of 63-66%, and precision of 44-49% (*Fig. 4B*). For CircleHunter, we report accuracies in the range of 70-76% and precision in the range of 40-48% (*Fig. 4B*). We report F1 and Matthew’s correlation coefficient (MCC) scores of ∼0.92 and ∼0.3 for AmpliconArchitect and ∼0.87 and ∼0.25 for CircleHunter. EcDNA predictions for both algorithms remain slightly underpowered for AmpliconArchitect, in that there are only 77 TP samples, when considering cell lines that have both WGS and karyotype annotations (*Fig. 4C*). For CircleHunter, only 25 TP samples have both ATAC-seq data and karyotype annotations (*Fig. 4D*). Thus, additional data will be necessary to increase the power of these computations.

**Figure 4.**
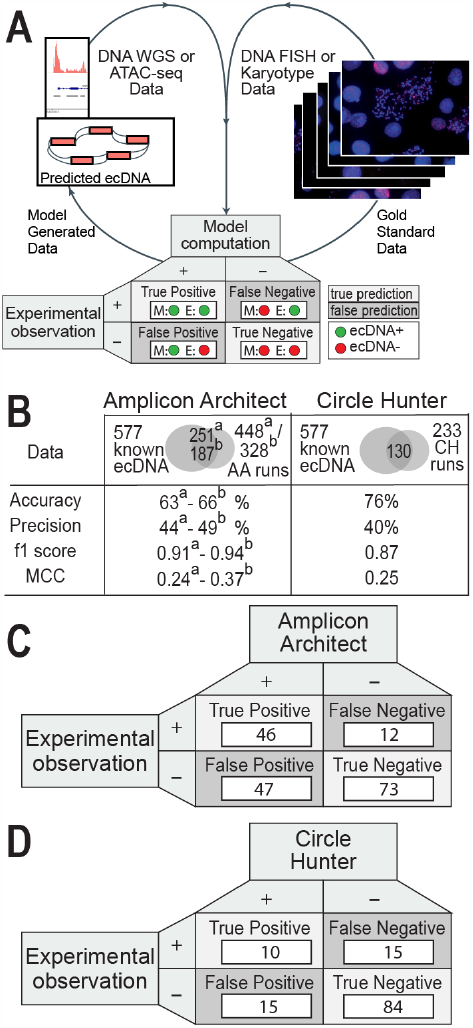
Comparison of computational predictions of ecDNA to karyotype annotations. **A**. Confusion matrices for two software programs that predict ecDNA were generated after assessing whether predictions aligned with karyotype-based ecDNA annotations. True positives (TP) refer to cell lines that are predicted to express ecDNA and are confirmed by karyotype whereas true negatives (TN) are cell lines that are not predicted to express ecDNA and karyotypes do not report ecDNA expression. False positives (FP) refer to cell lines that are computationally predicted to express ecDNA, but ecDNA expression was not observed experimentally. False negatives are cell lines that are computationally predicted to not express ecDNA but ecDNA expression was seen experimentally. **B**. Confusion matrix for AmpliconArchitect, in which accuracies are reported for 448 runs, which used default parameters, and 328 predictions from Amplicon Repository. **C**. Confusion matrix for CircleHunter, in which accuracies are reported from 233 runs, which used default parameters. **D**. A summary of accuracies, precision scores, f1 scores and MCC metrics for AmpliconArchitect and CircleHunter.

### Machine learning of karyotype features leads to high accuracy ecDNA and HSR prediction

A novel application of CytoCellDB is using karyotype information to predict ecDNA expression. Using a supervised learning approach, we demonstrate that we can predict ecDNA+ cell lines with high accuracy (AUC = 0.92). Furthermore, we developed a model capable of distinguishing between ecDNA+ and HSR+, and distinguishing between these types of gene amplifications with high accuracy (AUC = 0.88). These models open new avenues for identifying co-occurring genomic changes and chromosomal aberrations that may help understand ecDNA biogenesis and the relationships between ecDNA and genome instability.

The 400% increase in classifications enables the training of machine learning models with substantially more predictive power than any other ecDNA prediction model. We performed feature engineering of karyotype data for 577 cell lines by collecting 237 independent features from CytoCellDB’s manually curated karyotype data (*Fig 5A*). These features consist of polyploidy characteristics, chromosome losses and gains, marker chromosome characteristics, structural variation details and XY/chromosome complement details. As part of the feature engineering process, we transformed categorical columns (e.g., polyploidy characteristics) into numeric representations. In addition, we handled missing data in ploidy characteristics features by imputing them with ranges or means corresponding to their ploidy classification. The subset of cell lines with high coverage of these 237 features consisted of 94 ecDNA+/HSR-, 36 HSR+/ecDNA- and 399 ecDNA-/HSR-cell lines.

**Figure 5.**
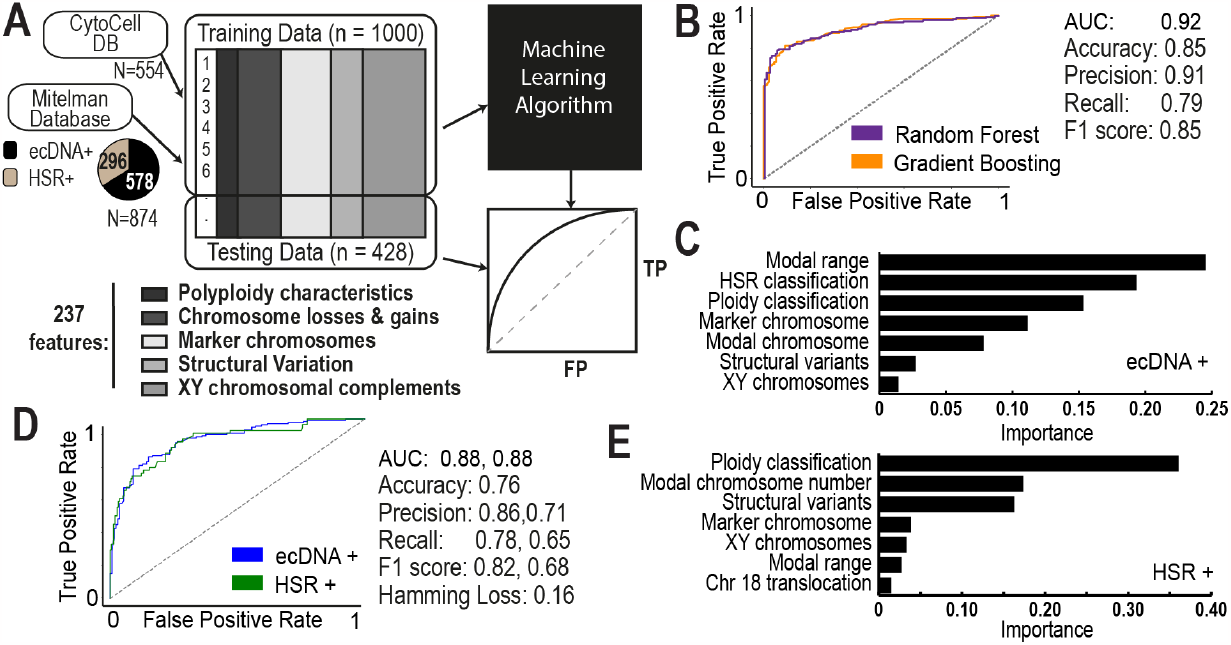
Machine Learning of Karyotype Data Predicts ecDNA and HSRs. **A**. A machine learning-based approach that takes in training karyotype data from CytoCellDB and the Mitelman database. Feature extraction identified 237 unique and independent features that were used to predict the expression of either ecDNA or HSRs. The training data was split so that 80% of the data was used for training and 20% of the data was used for testing. The final output was an analytic assessment of how well karyotype features predicted ecDNA versus HSRs. **B**. ROC curves of the candidate models indicated that both random forest and gradient boosting models performed reasonably well for ecDNA+ prediction, with AUC=0.92. **C**. Karyotype features with the highest importance during ecDNA+ prediction. **D**. ROC curves of the candidate models indicated that both random forest and gradient boosting models performed reasonably well for ecDNA/HSR+ prediction, with AUC=0.88. **E**. Karyotype features with the highest importance during HSR/ecDNA+ prediction.

To balance ecDNA+, HSR+ and ecDNA-/HSR-classes, we included 549 additional records of ecDNA+ cells and 266 HSR+ cells from the MitelMan database^16^. We performed uniform feature scaling and randomly partitioned the data into 80% for training and 20% for testing models. We trained various models on our data to compare performance, which included decision trees, support vector machines, random forests, gradient boosting, and bagging and stacking ensemble learning. The comparison of performance metrics revealed that the top-performing models were a Random Forest model and a Gradient Boosting model (*Fig 5B*). We found that both models performed exceptionally well, with AUC=0.92, accuracies of 85%, and f1 score of 85%. The top features that distinguish ecDNA+ and ecDNA-cells include modal chromosome range, HSR classification, ploidy classification, marker chromosome number, modal chromosome number, the total number of structural variants and the XY complement (*Fig 5C*).

Currently, there are no existing frameworks that can predict whether amplifications occur in ecDNA or in homogeneous staining regions (HSRs). This proves to be incredibly challenging because HSRs amplify genes in a similar manner to ecDNA, except they are in chromosomal DNA instead of extrachromosomal DNA. Extending our random forest model and gradient boosting model for binary ecDNA+ and ecDNA-classification to their multi-output classification model counterparts, we trained a multi-output random forest model and multi-output gradient boosting model with the similar approach using our set of 237 features to predict ecDNA+ and HSR+ cells. Our multi-output gradient boosting model outperformed the multi-output random forest model achieving reasonable accuracy predictions with prediction accuracy of 76%, an AUC of 0.88 for HSR and 0.83 for ecDNA, and f1 score of 82% for ecDNA, and 68% for HSR (*Fig. 5D*). The features with highest importance consisted of ploidy classification, modal chromosome number, total structural variance, marker chromosome details, XY complement, modal chromosome range and translocations involving chromosome 18.

### Pairwise Relationships Between Copy Number Variation and RNA Predicts ecDNA+ Cell Lines

RNA expression data has not yet been used to predict ecDNA or HSR in cell lines. However, when coupled with copy number variation data (CNV), it can provide useful insights into which genes are amplified and highly expressed by ecDNA. To test these relationships, we mapped CytoCellDB to CNV data and transcriptomics (RNA-seq) data from CCLE/DepMap. Several global patterns stand out when comparing pairwise relationships between CNV and RNA on a per-gene basis across ecDNA+, HSR+ and ecDNA-cell lines (*Fig. 6A*). One common motif is a clustering at the origin of the majority of cell lines with an “island” of cell lines that cluster in the upper right quadrant (high CNV, high RNA); this “island” of cell lines tends to be enriched in ecDNA+ cell lines for certain genes (e.g. p<8.7 e-4 using a chi squared test for cell lines amplifying MYC on ecDNA). We were interested in developing a systematic approach that uses these pairwise relationships to identify cell lines that express ecDNA.

**Figure 6.**
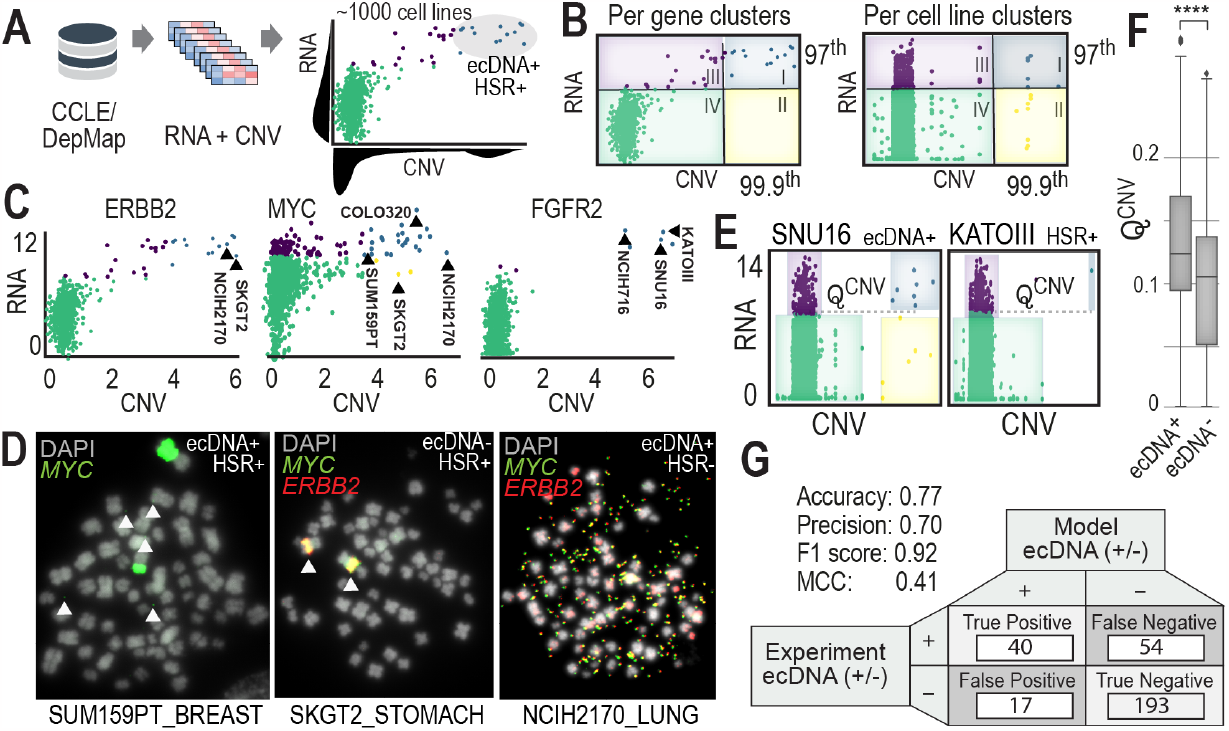
Multi-omics Analysis Finds Key Functional Relationships in ecDNA cell lines. **A**. Integration of bulk RNA-seq data with copy number variation data (derived from WGS) was taken from CCLE/DepMap. Pairwise analysis across genes and cell lines was performed and clustering analytics identified key patterns in cell lines expressing ecDNA. **B**. Clustering across all cell lines for a given gene (left) versus clustering across all genes for a given cell line (right). Clusters were assigned groups, based on quantile-based thresholding. **C**. Similar pairwise patterns in RNA vs CNV are seen for specific genes, such as ERBB2, MYC and FGFR2. Cell lines in cluster I (blue) have a high likelihood of expressing ecDNA or HSRs. This means that high copy number counts (amplification via ecDNA or HSR) is coupled with high expression of the same genes, indicating a functional relationship exists. **D**. Several cell lines with unknown ecDNA annotations were validated experimentally with DNA FISH. In two out of the three cases, genes were amplified on ecDNA and in one case on HSR. **E**. A parameter Q^CNV^ was derived to functionally differentiate genes amplified and expressed from ecDNA versus from HSR. **F**. Computing Q^CNV^ based on copy number data indicates that there is a difference in the number of genes amplified and the frequency at which they are amplified in the population that separates ecDNA+ cells from ecDNA-cells. **G**. Building an integrative predictive model for ecDNA expression considered functional clustering and predictions from AmpliconArchitect and CircleHunter. A confusion matrix reports the accuracy and precision metrics.

We looked at pairwise relationships of CNV and RNA on a: (i) per-gene level (across cell lines) and (ii) per cell line-level (across genes). We divided these pairwise plots into four quadrants using a quantile-based clustering approach (*Fig. 6B*, see Methods). We applied a genome-wide search and found 313 genes among 138 cell lines that shared the “island” motif. The genes most commonly observed in quadrant I are commonly amplified oncogenes, MYC-N (chr 2), MYC (chr 8), MDM2 (chr 12), CDK4 (chr 12), MIEN1 (chr 17), DDX1 (chr 2), FGFR2 (chr 10), and ERBB1 (chr 17). Examples of pairwise plots for ERBB2, MYC and FGFR2 are shown in *Fig. 6C*, where specific cell lines that express ecDNA or HSRs are labeled. For example, cell lines known to amplify MYC on ecDNA are COLO320DM^35^, NCIH716^36^, SNU16^37^ and those that amplify FGFR2 (fibroblast growth factor receptor 2) on HSRs include KATOIII^37^. A full list of genes and cell lines that follow this motif are included in Supplementary materials (Supp Table 1 & 2).

We identified three cell lines (NCIH2170, SKGT2, and SUM159PT) that were classified as quadrant I, which amplify ERBB2, MYC or both ERBB2 and MYC but do not have karyotype or cytogenetic data to confirm whether DNA was amplified on ecDNA or HSRs. We performed DNA FISH and G-banding experiments on these cell lines and found that two amplify DNA on ecDNA (SUM159PT and NCIH2170) and one amplifies DNA in HSRs (SKGT2). Our karyotype analyses indicated that SUM159PT had variable amounts of ecDNA that amplified MYC, ranging from 2-12 per cell. NCIH2170 had ecDNA in every cell, ranging from 100-1000 ecDNA per cell, that co-amplifies MYC and ERBB2. SKGT2 has 2-3 HSRs per cell that co-amplify MYC and ERBB2. Contrary to the hypothesis that ecDNA’s unique circular structure increases chromatin accessibility for high gene expression, pairwise RNA/CNV profiles for NCIH2170 and SKGT2 suggest that these cells achieve the same level of oncogene amplification and expression, regardless of whether the DNA is amplified on HSR or ecDNA.

Another global pattern that emerged from our analysis is that the number of highly amplified and highly expressed genes was distinct between ecDNA+, ecDNA- and HSR+ cell lines. We designed a parameter, Q^CNV^, to identify cell lines that had higher numbers of highly amplified and highly expressed genes (Fig 6E; Supp Fig. 2 and Methods). We computed Q^CNV^ across all genes and cell lines and found a significant difference between all groups, where ecDNA+ was significantly different compared to ecDNA-(Fig. 6F) and HSR+ (p<1e-4 using a Wilcoxon Rank Sum Test for two independent samples). We used Q^CNV^ as well as predictions from AmpliconArchitect and CircleHunter to improve the overall accuracy of predicting ecDNA to 77%, with a precision of 70%, F1 score of 0.92 and MCC of 0.41. These findings demonstrate the value in integrating across multiple omics data types to predict ecDNA.

## Conclusions

CytoCellDB is the first database to enable queries on chromosomal aberrations, including ecDNA, in commonly used cancer cell lines. In addition, it provides predictions of ecDNA and chromothripsis from computational algorithms, ShatterSeek^23^, AmpliconArchitect^13^ and CircleHunter^14^. Cell lines within CytoCellDB have been extensively characterized by CCLE/DepMap, which provides interoperable alignment across databases and access to a wide array of multi-omics data. Pairing CytoCellDB with the next generation of omics data-such as single cell datasets-will provide a compelling avenue for studying ecDNA at scale.

CytoCellDB provides new inroads for the field of cytogenomics, which merges cytogenetics research with genomics to study how chromosomal aberrations impact the genome at large. In total, it aggregates cytogenetics and karyotype data for 577 cell lines, which is a 400% increase in ecDNA annotations compared to what is currently known. Having a resource that brings all of this data together in one place will significantly advance research on ecDNA by providing new possibilities to study the impact of ecDNA on a global population scale, identifying new model systems that express ecDNA, and identifying tiered, publicly available multi-omics datasets that can be analyzed in the context of ecDNA. To increase accessibility of this resource, various karyotype features can be queried by users through an online database to link ecDNA expression, other chromosomal aberrations, and publicly available multi-omics data.

We demonstrate the utility of CytoCellDB through three applications. First, we used CytoCellDB to characterize the functional differences between ecDNA+ and ecDNA-cells in terms of gene expression, gene dependency and drug sensitivity differences. Second, karyotype annotations within CytoCellDB were used as features in machine learning models to predict ecDNA+, HSR+ and ecDNA-cell lines. Our models show that karyotype features faithfully predict ecDNA expression with accuracy as high as 85% and HSR expression with accuracy as high as 76%. Third, we predict ecDNA+, HSR+ and ecDNA-cell lines using paired copy number variation and RNA expression data. Our findings present preliminary, yet compelling, support for the potential of CytoCellDB to complement current prediction algorithms to increase their power, accuracy and precision. Future efforts to integrate functional omics data types and drug screens will advance our understanding of the broad molecular phenotypes that differentiate ecDNA+, HSR+ and ecDNA-cells.

CytoCellDB has limitations which are important to keep in mind. First, it is well known that cancer cell lines have unstable genomes which means karyotype data deposited in vendor databases or from past literature may not be up to date. Second, karyotype records were collected from different sources, which means some are higher quality than others. While true positives (TP) are less subjective and more straightforward to annotate, true negatives (TN) are harder to annotate because some ecDNA-cell line karyotypes definitively report the absence of ecDNA/HSR while others do not report ecDNA or HSRs. To address these challenges, karyotype records were classified as high, medium or low confidence, depending on the level of detail provided. Only the high and medium confidence records were used in analyses. While it would be extremely beneficial to have a single experimental platform with uniform conditions to characterize karyotype, performing cytogenetics experiments on 577 cell lines is currently cost prohibitive and technically challenging. Thus, CytoCellDB serves as a starting point to provide any available karyotype data on commonly used cancer cell lines. A major benefit of using CytoCellDB is comparing across multiple lines of evidence (experimental and computational) to determine the likelihood of a cell line expressing ecDNA or HSRs.

CytoCellDB provides a framework for exploring functional relationships in cell lines that express ecDNA and for assessing specific and genome-wide impacts of ecDNA expression. Integrated frameworks like CytoCellDB enable understanding of how ecDNA or HSRs influence the expression of key oncogenes or proteins, and how downstream network-level changes influence responses to drugs or CRISPR-mediated knock-down perturbations. CytoCellDB will aid in translating biomedical knowledge, from large, population-scale repositories like CCLE/DepMap to the discovery of novel therapeutics, targets, and clinical biomarkers that identify ecDNA in patient samples. Future efforts are likely to extend to precision medicine applications where patient tumor data can be analyzed and matched to drug therapies that target ecDNA-driven cancers. CytoCellDB is available online at (http://CytoCellDB.unc.edu).

## Methods

### Karyotype Mining

Karyotype data on CCLE/DepMap cancer cell lines was collected in two arms: (i) manual literature curation; (ii) direct algorithmic mining of cell line vendor databases. In the first arm, key words such as “double minutes,” “DMIN,” “metaphase cells,” “DNA FISH” along with cancer cell line identifers were used to search literature for existing manuscripts providing karyotype data and ecDNA classification. In total, 72 manuscripts were found that provided annotations on 157 cell lines. In the second arm, we designed a python script, using the python package BeautifulSoup^38^, to scrape the websites from two main cell line vendors: atcc.org and dsmz.de. A total of 643 unstructured karyotype records were linked to 577 cell lines using cellosaurus^39^. Karyotypes were manually curated by searching for the most common features typically detailed in karyotypes. These include modal chromosome number, modal chromosome range, ploidy classification, percent polyploidy, XY chromosome complement, chromosome losses or gains, marker chromosomes, structural variation, homogenous staining regions, double minute chromosomes. Where double minutes were observed and noted, we annotated these cell lines as having ecDNA. Where homogenous staining regions were observed and noted, we annotated these cell lines as having HSRs. For cases that did not report ecDNA or HSR, or for cases that reported that there were no ecDNA or HSRs observed, we annotated these cell lines as not having ecDNA or HSRs. Several cell lines had minimal information recorded for karyotypes. For example, only the ploidy was given. In these cases, we did not consider these as “negative” for ecDNA or HSRs.

### Machine Learning Analysis

Extracted karyotype data was engineered into 237 features used for machine learning analysis (see Supp File 1). Of the 554 CCLE/DepMap cell lines with detailed karyotype data, 108 were designated as ecDNA+ and 435 were designated with ecDNA-. We found that the imbalancing of these classes led to poor model results. To overcome this challenge, we integrated data from the Mitelman Database, which houses karyotype records on over 16,000 cell lines and tumor samples. We added 549 records for samples with ecDNA but not HSRs and 266 samples with HSRs but not ecDNA. In total, this amounted to 1428 total karyotypes (with 705 designated as ecDNA- and 712 designated as ecDNA+ and 11 designated as probable ecDNA+) used for machine learning analysis.

We trained various models on the dataset, compared their performance metrics, and selected two top-performing models: random forest and gradient boosting classification models. Additionally, we used a decision tree-based approach to determine which features generally separate ecDNA+, HSR+ and ecDNA-/HSR-cell lines. The effects of these features were measured using a Mann–Whitney-U test to determine the P-value of the separations. The outcome of such an analysis pipeline is to predict which cell lines are likely to have ecDNA/HSRs and understand what other karyotype features go hand in hand with extrachromosomal amplification.

Each dataset was randomly split, with 80% of the data used to train the machine learning models, and the remaining 20% of the data used to validate and test the model prediction results. The entire workflow for this project has been compiled into a series of user-friendly, scalable IPython notebooks, located at https://github.com/Brunk-Lab/CytoCellDB.

### Data Collection

WGS data for 468 cell lines and ATAC-seq data for 235 cell lines in the NIH Sequence Read Archive (SRA) was found using pysradb^40^ and was downloaded using SRA Toolkit. Copy number, bulk RNA-seq and CRISPR gene effect data was downloaded from the DepMap portal (depmap.org) from version 23Q2. Drug sensitivity data was downloaded from CTRP, GDSC1 and GDSC2. Metadata from DepMap / CCLE was obtained from version 21Q2 by cross referencing data availability for samples across each data type. Additional metadata for matching cell line identifiers across different databases was obtained from Cellosaurus^39^.

Karyotype data was downloaded from the Mitelman Database^16^ by accessing the data directly through the website (https://mitelmandatabase.isb-cgc.org/) and selecting “Cases Cytogenetic” and inputting “dmin” under “Abnormality.” Doing so, we retrieved 578 entries, using the version of the database that was last updated on October 16, 2023. We followed the same protocol to access entries with homogeneous staining regions (HSRs) by inputting “HSR” into “Abnormality.” This provided 278 entries (See Supp File 1).

ATAC-seq data was retrieved from the SRA database for 235 cell lines. After retrieving the SRA metadata, manual filtering was performed to remove genetically modified cell lines (i.e, parental cells genetically modified for KO, KD or gene overexpression) or under drug treatments from the dataset. All prediction experiments considered only control and/or vehicle treatments.

### Chromothripsis Prediction

We ran the software ShatterSeek on 184 cell lines using structural variation and copy number data downloaded from the DepMap portal and made prediction calls with high and low confidence based on the criteria recommended by the software^23^.

### Prediction of ecDNA using available softwares

AmpliconArchitect. The Amplicon Suite Pipeline was used to run both Amplicon Architect and Amplicon Classifier using WGS reads as FASTQ inputs aligned to hg38. Default parameters for the pipeline were used. No distinction was made between samples scoring above or below Amplicon Architect’s data quality threshold of a 95% BAM file properly paired rate. 447 of the 471 runs completed without errors.

CircleHunter. Corresponding SRA FASTQ files were downloaded using SRA Toolkit and CircleHunter was run with default parameters. For cell lines with multiple ATAC-seq samples, EcDNA detection was considered positive if at least one of the CircleHunter runs detected ecDNA.

### Power Analyses

We estimated the sample size needed to predict ecDNA at accuracies of 80% or higher using statsmodels^41^, by setting alpha to 0.05, beta to 0.80 and effect size to 0.4. The effect size was estimated using Cohen’s h for Proportions^34^ which states:

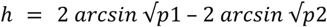

Where p1 and p2 represent ecDNA+ and ecDNA-populations. The proportions, p1 and p2, were estimated to be 0.4 and 0.6, respectively, which is consistently seen in several publications^2,15^. The effect size was computed to be 0.403, which is considered to be a medium effect size. Using these parameters, the sample size of 97 TP and 97 TN were determined to be sufficient.

### Integrative Functional Analyses

We compared the performance of manual clustering, density based clustering, k-means clustering, and gaussian mixture clustering. We found that density based clustering was the best non-biased based clustering method that worked well on pairwise RNA versus CNV data scatter plots. However, we ultimately chose to use manual clustering, using a quantile-based clustering method because this gave us the most control over the data. We generated clusters of data points based on the combination of their copy number values and RNA expression values. For example, cell lines or genes with genes with copy number (log2 relative to ploidy) > 3 and expression levels (log2 of TPM+1) > 7 were considered to be in quadrant I. Cell lines or genes with copy number (log2 relative to ploidy) < 3 and expression levels (log2 of TPM+1) > 7 were considered to be in quadrant II. Cell lines or genes with copy number (log2 relative to ploidy) < 3 and expression levels (log2 of TPM+1) > 7 were considered to be in quadrant III. Cell lines or genes with copy number (log2 relative to ploidy) < 3 and expression levels (log2 of TPM+1) < 7 were considered to be in quadrant IV.

We developed a parameter, Q, to describe the separation between genes likely to be amplified on ecDNA from genes (not amplified) on chromosomal DNA using the following equation:

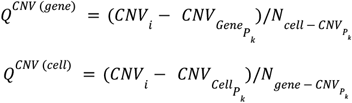

Q^CNV (gene)^ represents the difference between a gene’s copy number and the gene’s copy number value at the 99.9th percentile (P_k_ = 99.9) across the population of cell lines, divided by the total number of cell lines that have genes with copy number values at or above the 99.9th percentile.

Q^CNV (cell)^ represents the difference between a gene’s copy number and the copy number value at the 99th percentile (P_k_ = 99.9) across the genes within a given cell line, divided by the total number of genes within that cell line that have copy number values at or above the 99.9th percentile.

We integrated multiple features to predict ecDNA. The predicted set of cell lines were selected based on the following criteria:

1. Q^CNV (gene)^ > ***μ*** (Q^CNV (gene)^) +1σ (Q^CNV (gene)^) && Q^CNV (cell)^ > ***μ*** (Q^CNV (cell)^) +1σ (^QCNV (cell)^)
2. Cell lines are labeled as “quadrant I”
3. Cell lines are predicted to express ecDNA using AmpliconArchitect or Circle Hunter

### Primary Cell Culture

NCIH2170 and SKGT2 cells were purchased from ATCC and were grown in RPMI media (Gibco) supplemented with 10% heat-inactivated fetal bovine serum (Gibco) in a humidified incubator with 5% CO_2_. SUM159PT cells were given as a gift from Gary Johnson’s laboratory (UNC Chapel Hill) and were STR verified and checked for mycoplasma. SUM159PT cells were grown in Ham’s F-12 media supplemented with 5% heat inactivated fetal bovine serum along with 10mM HEPES, 1ug/ml Hydrocortisone, and 5ug/ml Insulin. Cells were harvested with 0.25% Trypsin in DPBS (Gibco). Viable cells were counted using a Countess 3 (Invitrogen) counter and trypan blue (Invitrogen, T10282). Cells were collected for metaphase and karyotype (G-banding) experiments within 3 passages.

### Metaphase Analyses

Cells were arrested at metaphase by overnight colcemid treatment (12-20h) at 0.1 μg/mL (10 μg/mL Colcemid Solution, FUJIFILM Irvine Scientific) in cell culture media, when cells were ∼70% confluent. Cells were harvested per standard cell culture procedure with minor alterations that trypsinization is quenched by a mixture of old colcemid-spiked media and PBS wash, to maximize the yield for semi-/adherent cells. Harvested cells were resuspended in 1 mL of 1x PBS by pipetting and transferred to a 1.5 mL microcentrifuge tube for centrifugation at 5000 rpm for 2 minutes. The supernatant was aspirated and each pellet was incubated with 600 μL of pre-warmed 37°C 0.075M KCl (Gibco) and at 37°C for 15 minutes in a water/bead bath. To quench the reaction we prepare Carony’s fixative (3:1 methanol:glacial acetic acid) fresh and add it dropwise in the tube. Tubes were immediately centrifuged at 5000 rpm for 2 minutes. We left 100-200 μL of supernatant to suspend pellets followed by addition of 600 μL of fixative dropwise and resuspension. This step was repeated 3 times. Depending on cell counts, we added 0-1 mL of fixative dropwise to make a 5-6 million cells/mL suspension. This ensures the optimal cell density for discernible spacing of single-cell karyotypes when dropped and imaged on microscope slides.

### Cytogenetic Analyses

Cytogenetic analysis was performed on twenty-five G-banded metaphase spreads of the human cancer cell line NCIH2170. The chromosome count in twenty-three of the spreads analyzed ranged from 62 to 68 (near-triploid). Two near-hexaploid spreads of 126 and 129 chromosomes were also observed. The majority of the spreads displayed a sex chromosome complement composed of two X chromosomes and one Y chromosome. Two spreads displayed one apparently normal X chromosome, and one X chromosome with additional chromatin of unknown origin attached to its p arm. One spread displayed a single X chromosome. Every spread displayed multiple chromosomal aberrations with minor variation from spread to spread. Multiple copies of apparently normal chromosomes were displayed, in particular chromosomes 7 and 20. Frequently observed structural aberrations include two variants of a derivative of chromosome 1 with a p arm deletion substituted with chromatin of unknown origin; an isochromosome of the chromosome 13 q arm; additional chromatin of unknown origin attached to the p arm of one copy of chromosome 13; two copies of chromosome 14 with an interstitial duplication of the distal 14 q arm; approximately 8 to 100 double minutes per spread; and two to seven marker chromosomes. A marker is defined as “a structurally abnormal chromosome that cannot be unambiguously identified by conventional banding cytogenetics.

Cytogenetic analysis was performed on twenty-five G-banded metaphase spreads of the human cancer cell line SKGT2. The chromosome count in twenty-three of the spreads analyzed ranged from 51 to 56 chromosomes per spread. Two near-tetraploid spreads of 104 and 108 chromosomes were also observed. Every near-diploid spread displayed a sex chromosome complement composed of one apparently normal X chromosome and one derivative X chromosome with a partial p arm deletion substituted with chromatin of unknown origin. Every spread displayed multiple chromosomal aberrations with minor variation from spread to spread. Consistently observed aberrations include an interstitial deletion in the proximal p arm of one copy of chromosome 3; a partial deletion of the chromosome 4 q arm terminus; a q arm deletion of one copy of chromosome 6 ; a partial deletion of the q arm terminus of one copy of chromosome 8; additional chromatin of unknown origin attached to the q arm of the other copy of chromosome 8; two variants of a chromosome 9 p arm deletion; additional chromatin of unknown origin attached to the p arm of one copy of chromosome 9; two copies of chromosome 10 with additional chromatin of unknown origin attached to the q arm terminus; two copies of chromosome 11 with additional chromatin of unknown origin attached to the p arm terminus; an isochromosome of the chromosome 13 q arm; a Robertsonian translocation resulting from the fusion of chromosome 15 and 21 q arms; two copies of a derivative of chromosome 17; monosomy of chromosomes 1, 4, 21 and 22; polysomy of chromosomes 14 and 20; and six to ten marker chromosomes. A marker is defined as “a structurally abnormal chromosome that cannot be unambiguously identified by conventional banding cytogenetics.

## Data availability

The workflow for this project has been compiled into a series of user-friendly Jupyter notebooks available at: https://github.com/Brunk-Lab/CytoCellDB

## Acknowledgements

The authors acknowledge funding support from the IBM Junior Faculty Developmental Award from University of North Carolina at Chapel Hill as well as support from Gary Johnson for providing the SUM159PT cell line, and Prof. Paul Mischel, Dr. Kristen Turner, Prof. Vaneet Bafna, Dr. Sudhir Chowdhry and Josh Lange on the helpful discussions early on that made this work possible.

## Author Contributions

EB conceived and managed the research. EB led, designed, and conducted analyses. JF, ST, HY, SH, SG, YW, NS, KG, and AM conducted computational analyses. JC, DC, and CGF conducted experiments. EB and CGF oversaw experiments and analyses. EB wrote the manuscript. EB and ST created figures for the manuscript. JF created the online database. All authors read and approved of the manuscript.

